# Inhibitory Units: An Organizing Nidus for Feature-Selective Sub-Networks in Area V1

**DOI:** 10.1101/282988

**Authors:** Ganna Palagina, Jochen F. Meyer, Stelios M. Smirnakis

## Abstract

Sensory stimuli are encoded by the joint firing of neuronal groups composed of pyramidal cells and interneurons, rather than single isolated neurons (Uhlhaas et al, 2009, Buzsaki, 2010). However, the principles by which these groups are organized to encode information remain poorly understood. A leading hypothesis is that similarly tuned pyramidal cells that preferentially connect to each other may form multi-cellular encoding units yoked to a similar purpose. The existence of such groups would be reflected on the profile of spontaneous events observed in neocortical networks. We used 2-photon calcium imaging to study spontaneous population-burst events in layer 2/3 of mouse area V1 during postnatal maturation (postnatal day 8–52). Throughout the period examined both size and duration of spontaneously occurring population-bursts formed scale-free distributions obeying a power law. The same was true for the degree of “functional connectivity,” a measure of pairwise synchrony across cells. These observations are consistent with a hierarchical small-world-net architecture, characterized by groups of cells with high local connectivity (“small worlds”, cliques) connected to each other via a restricted number of “hub” cells” (Bonifazi et al., 2009, Sporns, 2011, Luce & Perry, 1949). To identify candidate “small world” groups we searched for cells whose calcium events had a consistent temporal relationship to events recorded from local inhibitory interneurons. This was guided by the intuition that groups of neurons whose synchronous firing represents a “temporally coherent computational unit” (or feature) ought to be inhibited together. This strategy allowed us to identify clusters of pyramidal neurons whose firing is temporally “linked” to one or more local interneurons. These “small-world” clusters did not remain static, during postnatal development: both cluster size and overlap with other clusters decreased over time as pyramidal neurons became progressively more selective, “linking” to fewer neighboring interneurons. Notably, pyramidal neurons in a cluster show higher tuning function similarity than expected with each other and with their “linked” interneurons. Our findings suggest that spontaneous population events in the visual cortex are shaped by “small-world” networks of pyramidal neurons that share functional properties and work in concert with one or more local interneurons. We argue that such groups represent a fundamental neocortical unit of computation at the population level.

## RESULTS

### Scale-Free Population Burst-Events in Layer 2/3 of Mouse Area V1 Reveal “Small-World” Network Architecture

We imaged spontaneous ongoing activity in the developing postnatal and juvenile-adult visual cortex of mice using 2-photon calcium imaging. OGB-1 dye was used to densely label neocortical neurons at early postnatal ages. 1-4 movies were acquired per field of view (FOV) at 3.23-11 Hz under light (0.6%) isoflurane anesthesia (∼11-40 minutes in total). FOVs contained on average ∼148 (78-292) neurons whose calcium signals were acquired simultaneously (Fig.1A&B). The onset of individual OBG calcium events was identified by thresholding the dF/F trace at +3 standard deviations from the level of noise in each cell (Fig. 1B). The threshold selected ensured that identified calcium events reflect burst spiking events with high likelihood: ∼77% of spike burst onsets (doublets and multiplets) were detected reliably (see Methods). For each cell, we assigned the number 1 to frames containing a calcium event onset, while frames with no event onsets were assigned the number zero. This yields a binary “eventogram” of firing event onsets across the whole imaging period. We then grouped events into spatio-temporal multi-neuronal bursts (Fig. 1B, bottom panel). The beginning of a burst was defined as a frame that contained at least one cell event onset following an event-free epoch. The burst continued as long as consecutive frames contained at least one calcium event onset in one of the FOV cells, and ended on encountering an event-free frame. All identified neurons generated spontaneous calcium events. We focus on population burst-events with number of participating cells (size) s >1. We detected on average ∼1200 spontaneous multi-neuronal burst-events per FOV, of which ∼55% had size >1 (burst-event rate ∼0.6-1 Hz). Burst-event rate decreased slightly during postnatal development (Fig. 1D), following the event rate decrease seen in individual cells (Fig.1C,D). At every postnatal age examined the distribution of burst-event sizes was scale-free and conformed to a power law (Fig. 1E). Scale-invariance suggests that firing events happen at all spatial scales up to the size of the studied FOV. The scaling factor (α), i.e. the slope of the linear log-log plot of burst-event probability vs size, characterizes the relative probability of occurrence of burst-events of different size. Scaling factors for burst-event sizes remain close (a∼1.4–1.97) throughout all ages examined (Fig.1E), from before eye opening (P8-P10) to juvenile adulthood (P35+). The fact that burst-event sizes obey a power law implies that individual neuronal events are functionally coupled (Eurich et al., 2002, Beggs & Plenz, 2003) throughout postnatal development.

**Figure 1.**
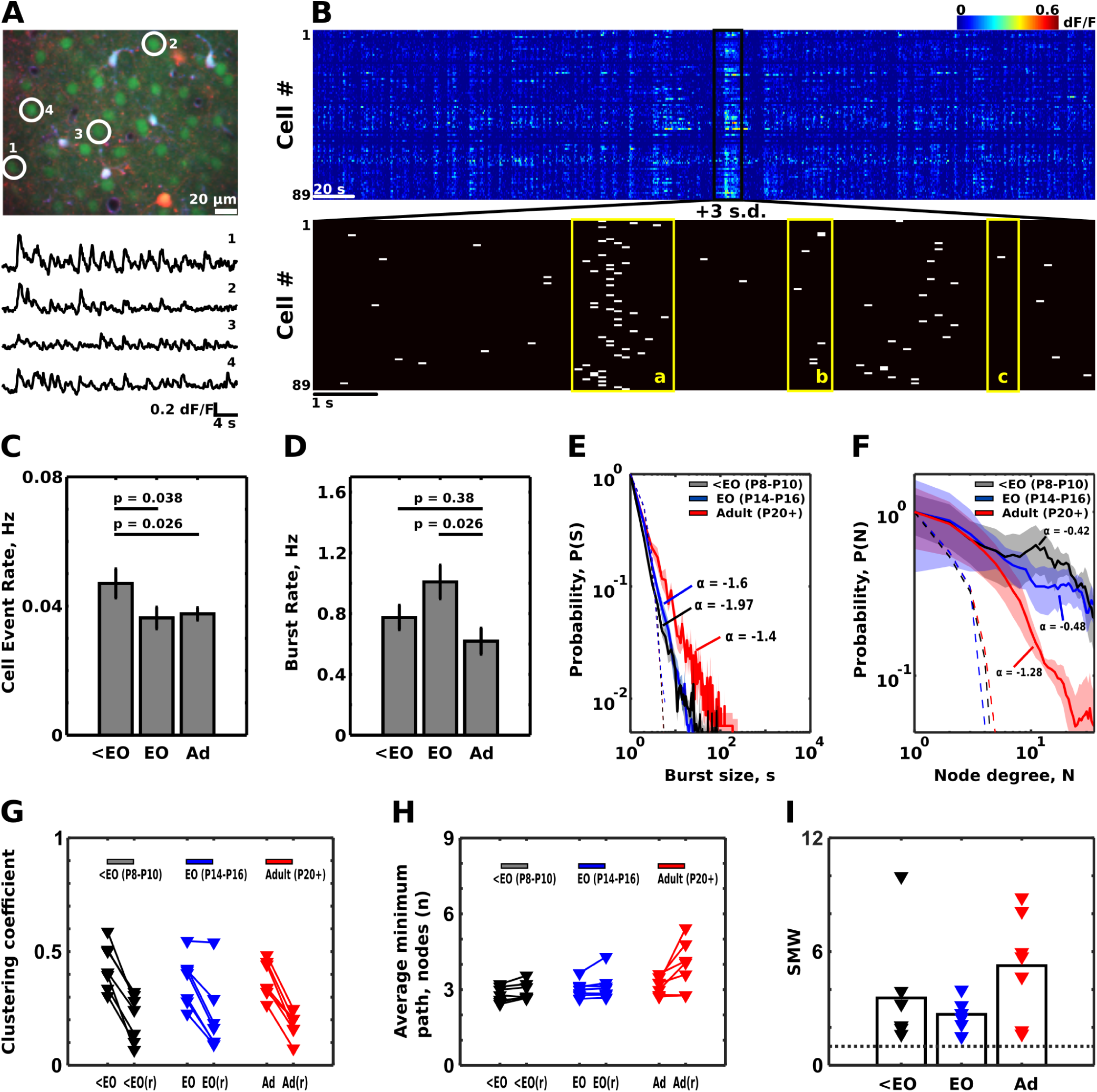
Spontaneous activity organizes as multi-neuronal population bursts. **A. Top panel:** FOV in the primary visual cortex stained with OGB-1 (green cell bodies). Red cells: Td- Tomato labeled Dlx5/6+ interneurons. Light blue: astrocytes (labeled by SR-101). **Bottom panel:** Example calcium dF/F signal timecourses from cells circled in A. **B. Top panel:** Calcium dF/F responses from a population of 89 cells. Dark outline: Sub-region expanded in the bottom panel. **Bottom panel:** population eventogram showing the onset of calcium responses (3 standard deviations above baseline activity) in each individual cell. Population bursts are characterized by their size and duration (see methods). The start of a population burst is a frame that contains at least one active cell preceded by an empty frame; the end is the frame that contains at least one active cell followed by an empty frame. Yellow boxes (a – c) outline example bursts. Box a: 46 cell, 1.4 sec long event. Box b: 7 cell, 0.46 sec. Box c: solitary event (size=1 cell). **C.** The rates of events in individual cells slightly decrease after eye opening (<EO vs Adult, p = 0.026; Wilcoxon ranksum test). **D.** The rate of population bursts with size >1 also somewht decreases after eye opening (EO vs Adult: p = 0.026). **E,F.** Properties of population bursts follow scale-free distributions (<EO, n=7, EO, n=7, Adult, n=7). **E.** The distribution of the event sizes obeys a power law (α = −1.4 to −1.97) at every developmental stage tested (<EO: P8-P10; EO: P12-P16; Adult: P35+). *<u>Solid lines</u>*: mean across individual FOV distributions; shading: SEM. *<u>Dotted lines</u>*: Control, poisson distribution matched for size. **F.** The distribution of node degrees, i.e. the number of significant functional links formed by each neuron (see Methods), are also scale-free at every developmental stage. However, now the power law changes slope (from −0.42 before eye opening to −1.28 in adulthood) reflecting a reduction in the number of functional links with postnatal maturation. *<u>Solid lines</u>*: mean across individual FOV distributions; shading: SEM. *<u>Dotted lines</u>*: Control, poisson distribution matched for node degree. **G-J.** V1 layer 2/3 V1 networks display small-world properties at every developmental stage. **G.** Clustering coefficients in L2/3 networks are larger then corresponding clustering coefficients in random networks with similar average number of links per node. <EO: before eye opening, EO: around eye opening, Ad: adult. <EO(r), EO(r), Ad(r) represent corresponding clustering coefficient values obtained from random networks with similar average number of connections per cell. These values were conservatively taken to be 3 standard deviations (99.7% cutoff) above the mean of a null distribution of clustering coefficient values obtained by randomly re-assigning existing connected pairs across the network nodes 5000 times (see Methods). **H.** Average minimal path length stays similar across ages and is slightly reduced compared to random networks with similar total number of nodes and average number of links per individual node. Conventions are similar to G. **J.** Small world coefficients (SMW), obtained by normalizing the real clustering coefficient and average minimum path length by those of corresponding randomly connected networks (see Methods) exceeds ∼1 (grey dotted line) at every age examined.

To further analyze the structure of synchronous events, we assigned a link between pairs of cells whose Pearson correlation coefficients reached a desired level of significance (p < 0.0001, see Methods). The number of links emanating from each cell measures the cell’s “degree of connectivity,” i.e. the number of neighboring units with above chance probability to fire in synchrony with that cell. The degree of “functional connectivity” across L2/3 cells also obeyed a power law (Fig. 1F), suggesting an underlying hierarchical small-world network architecture (Sporns, 2011). This architecture features “cliques” of functionally inter-connected cells forming sparse links with each other via relatively rare clique cross-connecting hub nodes (Sporns, 2011, Bassett & Bullmore, 2006). Hierarchical small-world nets are characterized by large average clustering coefficient to average minimal path length ratios compared to randomly connected nets (see Methods), i.e. by a “small world factor” (SMW) greater than 1 (Sporns, 2011) (Fig. 1G,H,I). V1 layer 2/3 has SMW >>1 (range 1.52-9.96) at every postnatal age examined (Fig.1I). Interestingly, the magnitude of the scaling factor for the degree of connectivity increased dramatically with postnatal maturation (from −0.42 to −1.28; Fig.1F), suggesting gradual functional link elimination and subsequent decrease in the size of highly connected “cliques” over time after eye opening (Fig. 1F). This is in line with spontaneous activity de-correlation during the first weeks of postnatal development (Golshani et al., 2009, Rochefort et al, 2009, Fig. 3C).

Cliques of neurons in the adult brain have been reported in the literature (Markov et al., 2013, Meunier et al, 2010, Sporns & Honey, 2006, Humphries et al., 2006) and are proposed to play an important role in cortical computations (Lago-Fernandes et al., 2000). We aimed to identify and characterize putative “small-world” cliques in L2/3 of mouse area V1 and investigate how they evolve during development. Our approach is guided by the intuition that cliques of functionally similar pyramidal cells whose synchronous firing represents a “temporally coherent computational unit” (or feature) ought to be inhibited together. Therefore, clique members are expected to form functional connections not just with each other, but also with the same set of local neighboring interneurons.

### Sub-Networks of Pyramidal Neurons Temporally Linked to Specific Interneurons

To identify candidate “small world” groups we searched for cells whose calcium events had a consistent temporal relationship to events recorded from local inhibitory interneurons. We used mice expressing Td-Tomato in a subset (∼ 65%) of layer 2/3 interneurons positive for the protein Dlx-5/6 (Madisen et al., 2010). For each interneuron, we identified the pyramidal cells that generated events with higher probability within the 600 ms window immediately preceding interneuronal firing events (Fig. 2A). This relatively large window reflects the build-up of recurrent network activity rather than purely monosynaptic connections, which are faster (see Extended Data Fig.1, Methods). Significance was estimated by random circular shuffling of calcium response onsets to build a null distribution of pyramidal cell event probability relative to interneuronal events (Fig. 2B; p = 0.0001 (0.0045 after correcting for multiple comparisons), and less than cutoff p-value of p=0.0011 (which corresponds to 0.05 after correction for multiple comparisons)). These pyramidal cells have the potential of activating the interneuron and then being themselves inhibited by the interneuron in the context of recurrent network activity. We refer to such cells as “*<u>partners</u>*“ of that interneuron. An interneuron with its associated pyramidal “partners” define a “clique” we call “interneuron pyramidal partner cluster“, or “IPP-cluster” (Fig. 2C-E). Nearly all (∼97%) interneurons had pyramidal partners inside the FOV. Interestingly, the average distance of IPP-cluster pyramidal members from the partner interneuron remained stable across the ages examined (Fig. 2H) when normalized appropriately by the increase in inter-neuronal distance expected during the course of postnatal development and FOV size (maximal distance between cells) (Fig. 2H). Note that IPP-cluster identification depends on *<u>functional</u>* connectivity between interneurons and pyramidal cells without implying anything about the underlying anatomical substrate.

**Figure 2.**
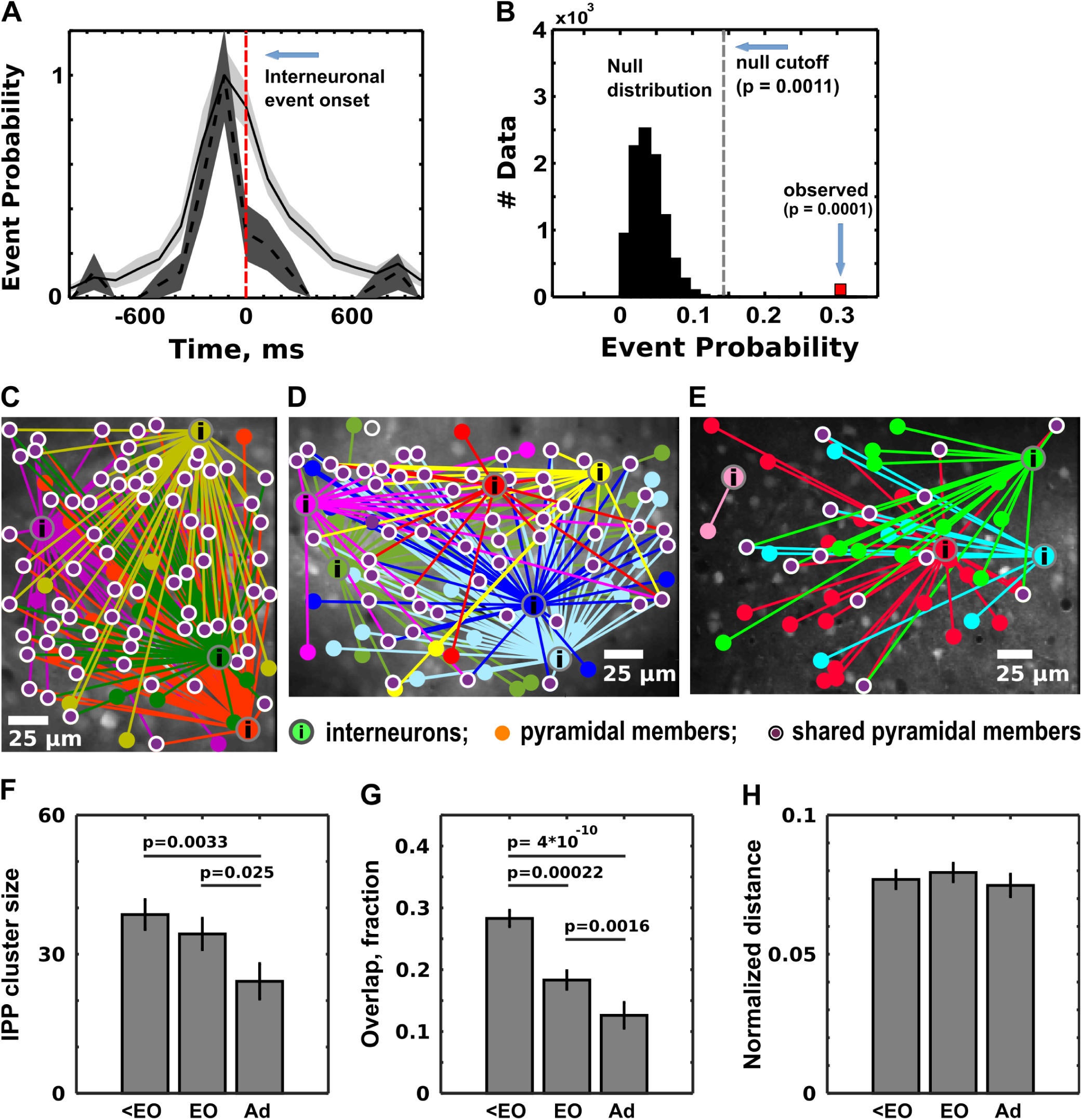
Interneuron Pyramidal Partner Clusters (IPP-clusters). Identifying pyramidal cells with increased probability to fire calcium events prior to the interneuron’s events. **A.** Normalized histogram of pyramidal cell events occurring [-1000, 1000] ms around the calcium response onsets of the interneuron. *Dotted black line*: example of a single pyramidal cell (mean over 65 trials). *Solid black line*: aggregate responses of all pyramidal cells in an FOV whose probability of firing was increased prior to the interneuron’s events. Filled patches: standard error of the mean across all trials. **B.** To assess significance, we first selected out pyramidal cells which had larger probability of events in the 600 ms period before the events of the interneuron compared to 600 ms period after the events of the interneuron. Next, the event trains of these pyramidal cells were randomly shuffled cell-by-cell, in circular fashion, to generate the 10000 point null distributi n of pyramidal cell event probability in 600 ms window after the events of the interneuron. Th significance of the difference for each prospective pyramidal partner could then be determined b comparing the pyramidal cell’s event probability prior to interneuron’s event relative to the null distribution (black histogram). The threshold p-value was set at 0.05 after correction for ultiple comparisons (in the example above this corresponds to initial uncorrected p-value of 0.0 11). Cells with p-values below the threshold were accepted as partners, denoting a link between th interneuron and the corresponding pyramidal cell (red bar, uncorrected p-value is 0.0001, corresp nding to 0.0045 after correction). C-E. Examples of IPP- clusters at different developmental p ints. C: P9, D: P15, E : P35.IP P-cluster members are color coded. i: interneurons, f***illed disks: p*** ramidal memrbse, circled disks: s hared pyramidal members. Bars: 25 μm. F. IPP-clus t er siz decreases over time during postnatal development (<EO: P8-P10, n = 45; EO: P12-P16, n = 41; Ad: P35 - P52, n = 38). G. IPP-cluster overlap also decreases over time. Conventions as in F. H. The averagei dstance of pyramidal IPP-cluster members from their partner interneuron, normalized for he expected increase in inter-neuronal distance over postnatal development, remains stable ith age (see Methods). Significance reported in F and G was determined by the Wilcoxon ranksu test.

### Evolution of IPP Clusters During Postnatal Development

The number of neurons per IPP-cluster as well as average overlap between IPP-clusters decreased significantly during postnatal development (Fig. 2F,G). Specifically, before eye opening (P8-P10) mean IPP-cluster size was ∼39 pyramidal neurons (n=45), around eye opening (P12-P16) ∼34 (n=41), while in juvenile adulthood (P35-P52) it dropped further to ∼24 cells (n=38) (Fig. 2F). Correspondingly, average overlap between the clusters was ∼28% before eye-opening, ∼18% around eye opening, and ∼13% in juvenile adulthood (Fig. 2G). Note that there was no systematic difference in the fraction of active cells per FOV across different groups and IPP-cluster size/overlap did not correlate with the number of active neurons in the FOV.

To measure the strength of pairwise functional connectivity within a cluster we used linear Pearson cross-correlation function over a ±600ms window *<u>after</u>* removing all [-600ms,600ms] periods centered around interneuronal responses to exclude periods in which interneurons can directly influence their pyramidal partners (or vice-versa; see Methods). Pairwise functional connectivity strength was greater between pyramidal IPP- cluster members versus IPP-cluster members and appropriately distance matched non members from the same FOV (Fig. 3C; see Methods). The difference in mean pairwise functional connectivity strength was significant already before eye-opening and remained significant through adulthood (Fig.3C). We computed the mean pairwise connectivity strength across all pyramidal members of an IPP-cluster and compared it to that derived from a null distribution generated by creating 10000 surrogate size-matched and distance-matched pyramidal cell groups randomly drawn from the FOV without replacement (see Methods; Fig. 3A,B). IPP-clusters with mean pairwise functional connectivity strength >99.7% of the null distribution’s values were conservatively deemed to have mean pairwise functional connectivity strength higher than chance. Remarkably, the majority of IPP- clusters exceeded this threshold; irrespective of postnatal age the fraction of such clusters exceeded 60% (Fig. 3D). *Note that at the threshold chosen only ∼1% of IPP-clusters were expected to reach significance by chance, confirming that IPP-cluster* represent well defined cliques of relatively densely inter-connected cells.

**Figure 3.**
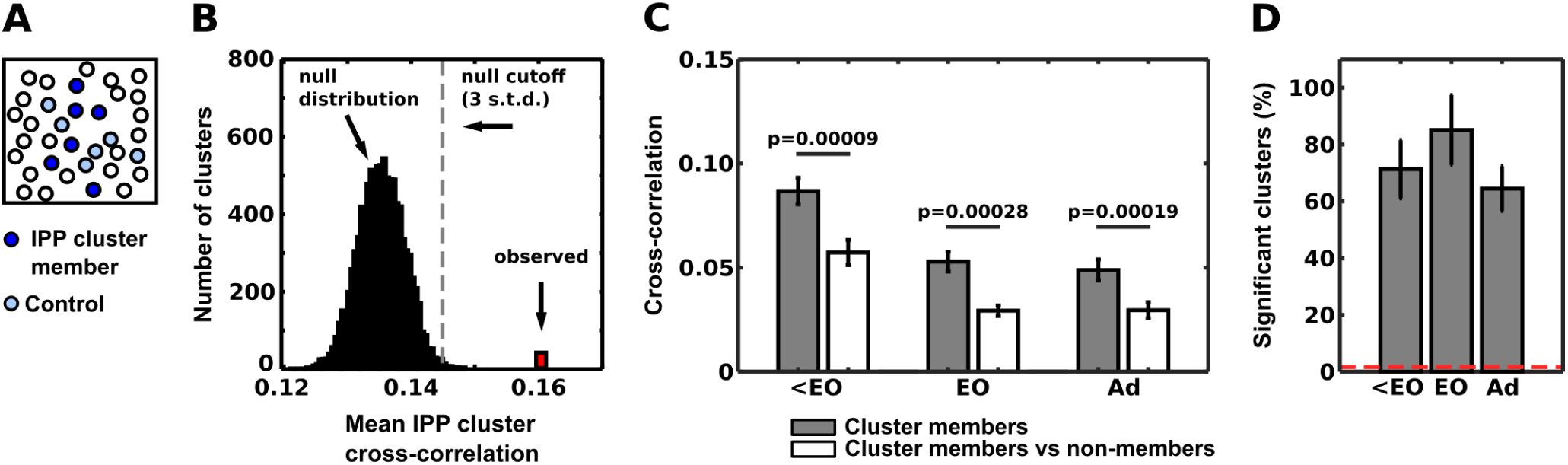
IPP-cluster members show higher pairwise cross-correlation strength. **A.** *<u>Dark blue:</u>* IPP cluster members. *Cyan:* size- and distance-matched control sample of randomly selected pyramidal cells from the same FOV. This control cluster has to satisfy the following requirements: i) its size corresponds to original cluster’s size, and ii) its members have to be on average at the same distance from the original IPP-cluster’s partner interneuron as the IPP-cluster’s pyramidal members (see Methods). **B.** Null distribution (black histogram) of mean pairwise Pearson cross-correlation values (600 ms window) computed between pyramidal cells, one of which belongs to the IPP-cluster, the other to a control cluster (see Methods). Ten thousand control clusters were randomly selected per IPP-cluster within the FOV. Significance required that the mean pairwise correlation coefficient of the IPP-cluster (red) be greater than 99.7% of the null distribution values. **C.** Pairwise cross correlation strength between IPP-cluster pyramidal partner members that belong to the same cluster (gray bars) versus pyramidal non-members, after appropriately controlling for pairwise distance (see Methods). Pairwise correlation strength decreases over time in agreement with the decorrelation that occurs in early postnatal development (Golshani et al., 2009, Rochefort et al., 2009). However, at all ages, average pairwise cross correlation strength is higher within versus across different IPP-clusters. P-values are obtained by Wilcoxon ranksum test. **D.** Percentage of IPP-clusters whose members have significantly higher mean cross correlation strength than control, using the criterion described in B. Chance level is ∼1% (red dotted line). The percentage of IPP-clusters with significantly higher mean pairwise cross correlation strength than control is already substantial before eye opening (∼74% for <EO: P8-P10, n = 44 clusters from 7 FOVs) and remains high (65-85%) after eye opening (EO: n=40 clusters from 7 FOVs, Adult: n = 36 clusters from 7 FOVs). Clusters consisting of the iterneuron and single connected pyramidal cell were excluded from analysis.

“Cluster-exclusive” pyramidal cells, i.e. cells that participate exclusively in a single IPP-cluster form fewer functional links than “shared” pyramidal cells (cells that participate in more than one IPP-cluster) at all ages examined (Fig.4A). However, both “cluster-exclusive” pyramidal cells and “shared” pyramidal cells retain fewer functional links over time and the number of functional links per pyramidal IPP-cluster member decreases over postnatal development (Fig. 4A). The fraction of “cluster-exclusive” pyramidal cells greatly increases after eye opening (Fig. 4C), while the fraction of “shared” pyramidal cells correspondingly decreases. As a result, by juvenile adulthood a pyramidal cell member of an IPP-cluster is on average functionally linked to ∼2 interneurons, down from ∼3.5 prior to eye opening (Fig. 4B). Along the same lines, the average number of clusters cross-linked by a “shared cell” decreases from ∼4 (before eye opening) to ∼3 (juvenile adulthood), (Extended Data Fig.4). These processes reflect the refinement of small-world network structure, with IPP- clusters becoming increasingly segregated, reflecting perhaps an increase in functional specialization (Gao et al., 2010).

**Figure 4.**
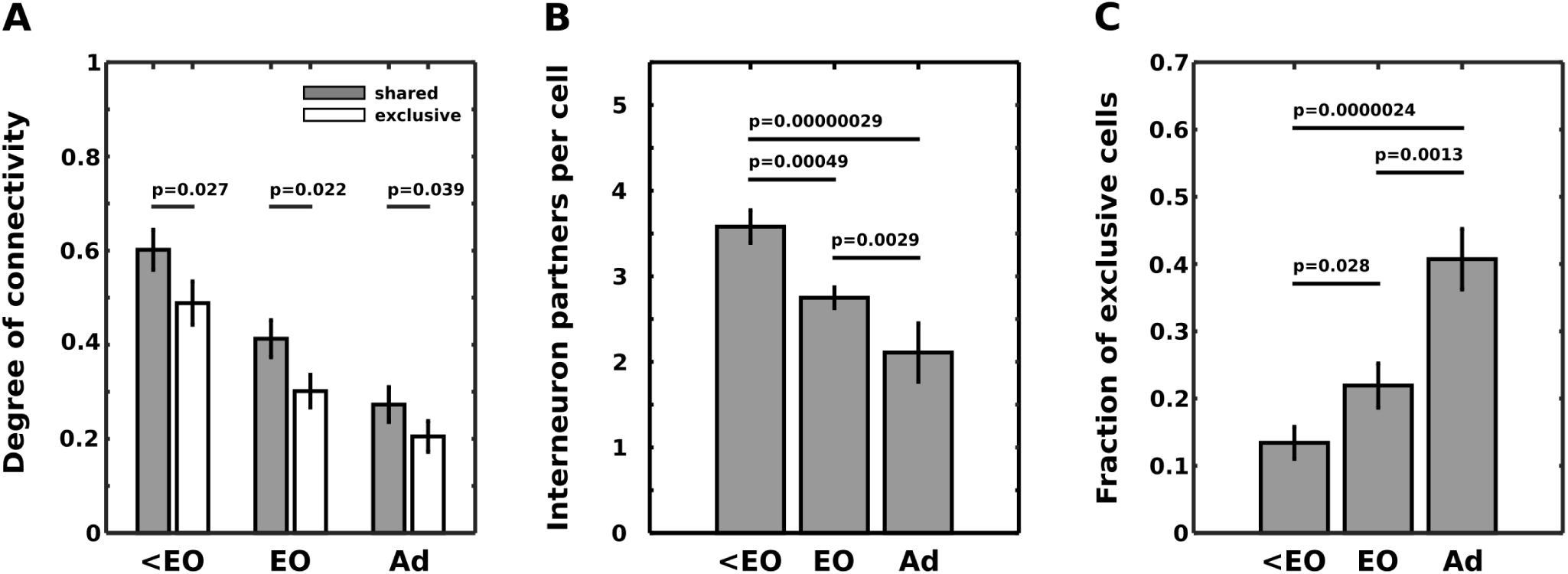
Refinement of IPP-cluster functional connections during development. **A.** The degree of functional connectivity (number of functional functional links per pyramidal cell, expressed as a fraction of total number of pyramidal cells in the FOV) decreases over the course of early development. Pyramidal cells participating in more then one IPP-cluster (“shared” or “hub” cells; gray bars) have higher degree than cells participating exclusively in one IPP cluster (“exclusive”; white bars) at all ages examined. On average the degree of connectivity for shared cells drops from 0.6±0.043 before eye opening (<EO, n=42 clusters from 7 FOVs) to 0.41±0.04 around eye opening (EO, n=35 clusters from 7 FOVs) to 0.27±0.04 in adulthood (P35+, n=33 clusters from 7 FOVs). Corresponding numbers for “exclusive” cells are 0.49±0.046 to 0.3±0.035 to 0.205±0.03 respectively. The degree difference between shared and exclusive cells is significant for every age group (ranksum test: p = 0.022 – 0.039, signed rank test: p = 5.3*10 ^-7^ – 1.4*10^-5^), even though both exclusive and shared cells lose significant number of connections after eye opening and with further development (ranksum test; shared cells: p(<EO vs EO) = 0.0088, p(EO vs adult) = 0.03; exclusive cells: p(<EO vs EO) = 0.0019, p(EO vs adult) = 0.0185). Note, that only clusters containing both exclusive and shared members were used in this analysis. **B.** The number of interneuron “partners” per pyramidal cell decreases on average during developmental refinement from ∼3.5 (<EO, n = 45 clusters, 7 FOVs), to ∼2.7 (EO, n = 41 clusters, 7 FOVs) to ∼2.1 interneurons / pyramidal cell (adult, n = 38 clusters, 7 FOVs). Statistical significance was determined by the Wilcoxon ranksum test across clusters. **C.** The fraction of pyramidal cells that participate exclusively in one IPP-cluster increases over time, from 0.13±0.02 (<EO, n = 45 clusters, 7 FOVs) to 0.22±0.03 (EO, n = 41 clusters, 7 FOVs) to 0.41±0.045 (Adult, n = 38 clusters, 7 FOVs).

Our findings argue strongly that IPP-clusters represent neuronal cliques existing as early as P8, whose structure undergoes marked reorganization during early postnatal development. Specifically, IPP-clusters decrease in size and become more disjoint as functional connections between their pyramidal members get pruned over time. Notably, the average number of IPP-clusters linked per “shared” (“hub”) pyramidal cell decreases over time, as the network acquires incrementally stronger “small-world” structure. So far, all analysis was performed on epochs of spontaneous activity. The question then arises whether IPP-cluster members share functional properties, and whether “shared” nodes cross-link IPP-clusters with similar functional properties.

### IPP-cluster members share physiological properties

We found that pyramidal members of IPP-clusters share functional properties with each other and with their partner interneurons. This analysis was performed in 4 juvenile adult animals (P42-P52, 4 FOVs), in which direction and orientation selectivity are mature (Rochefort et al., 2011). ∼52% (12/23) of visually responsive interneurons recorded in adult animals showed direction- and/or orientation-selectivity when tested with drifting gratings and plaid textures. Fig.5A,B illustrates an example of three direction tuned pyramidal IPP-cluster members linked to their tuned interneuron partner. Note that the peaks of the tuning functions align across the pyramidal cells and the interneuron (Fig.5C). Fig.5D-E summarizes population data across all IPP-clusters derived from oriented interneurons tested with moving gratings: ∼34% of pyramidal members share direction preference with their partner interneuron, ∼75% are within ±45°, and only ∼14% have preference orthogonal (at least 90 degrees difference) to that of their partner interneuron (Fig. 5D,E, p = 0.0049). This observation is not strictly specific to moving gratings but translates to different types of moving stimuli, such as moving textures (Extended Data Fig.2). Extended Data figures 2 and 3 demonstrates high tuning-curve similarity between interneurons and their pyramidal partners for a moving plaid with 120° cross-angle. Accordingly, population-tuning curves derived from the responses of all IPP-cluster members were highly correlated with the tuning curve of their ‘partner’ interneurons, across all conditions tested (see methods): median linear Pearson correlation of 0.58 (n = 12; gratings), 0.51 (n = 10; 120°-cross-angle plaid). The similarity in tuning between cluster members and partner interneuron is stimulus-independent even though the tuning itself is stimulus specific: a change in the visual stimulus causes similar changes in the tuning function of both the interneuron and its partner pyramidal IPP-cluster members (Extended Data Fig. 3).

**Figure 5.**
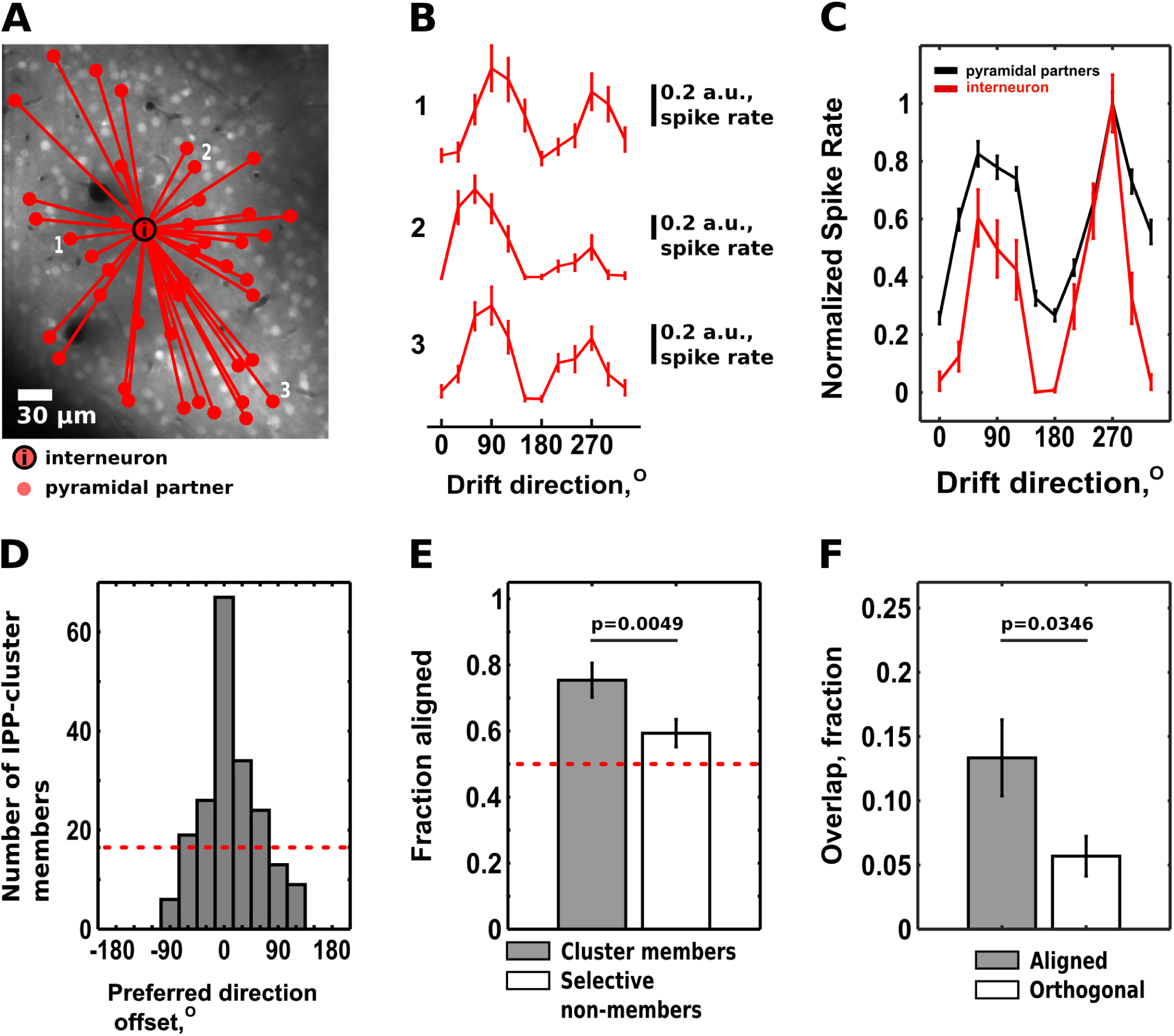
IPP-cluster members have similar functional properties. **A.** Example adult V1 L2/3 IPP-cluster with 45 pyramidal cell members. The partner interneuron is denoted by “**i**”. **B.** Example tuning functions to drifting gratings for cells 1-3 shown in A. **C.** Tuning function derived from the pyramidal members of the IPP-cluster shown in A (black; see methods) compared to the tuning function of the partner interneuron (red). Note that they share high similarity (median Pearson correlation coefficient ∼0.58 over n = 12 IPP clusters). **D.** Distribution of differences in the preferred direction of motion between tuned IPP-cluster pyramidal cell members and their partner interneuron, whose preferred direction is by convention set to zero (see methods). The majority of tuned IPP cluster members have the same preferred direction as their partner interneuron. ***<u>Red dotted line:</u>*** fraction expected by chance. **E. *<u>Gray Bar:</u>*** Fraction of IPP-cluster pyramidal cells whose preferred direction is within ±45° of the preferred directions of their partner interneuron. ***<u>White Bar:</u>*** Control (tuned pyramidal cells selected randomly in the vicinity of the interneuron). ***<u>Red dotted line:</u>*** Fraction expected by chance. The difference is significant (p = 0.0049, Wilcoxon signed rank test, n = 12 IPP-clusters). Data are expressed as mean±sem; cluster members: 0.754±0.052, oriented non-members: 0.59±0.042. **F.** IPP-clusters with similar (within±45°) tuning (“aligned”) show significantly larger pyramidal cell overlap compared to IPP-clusters with “orthogonal” (more than ±45°) preferences, which have very small overlap. ***<u>Gray Bar:</u>*** aligned clusters, 0.133±0.03 (n = 10 IPP-cluster pairs). ***<u>White Bar:</u>*** orthogonal clusters, 0.057±0.015 (n=16 IPP-cluster pairs). Statistical significance was determined by the Wilcoxon ranksum test.

We conclude that pyramidal members of IPP-clusters share tuning properties with each other and with their ‘partner’ interneuron. Note that visual tuning properties did not necessarily have to be *a-priori* similar across members of the same IPP-cluster, since IPP-clusters were defined by analyzing spontaneous patterns of activity in V1 L2/3 *in the absence of visual stimulation*. The fact that they are suggests that IPP-clusters have functional significance for the encoding of visual stimuli. In accordance with this, “shared” nodes preferentially link IPP- clusters with similar tuning properties. Fig. 5F shows that IPP-clusters with similar direction preference (within 45 degrees of each other) had on average ∼13% “shared” nodes, whereas IPP-clusters with orthogonal preference (difference > 45 degrees) shared only ∼6% of nodes (Fig. 5F, p = 0.0346). Therefore IPP-clusters with similar feature-preference retain stronger connectivity through “shared” high-degree nodes, while IPP-clusters with orthogonal tuning largely separate from each other in the adult neocortex.

## DISCUSSION

Understanding how neocortical neurons organize into functional groups that multiplex with each other to process information is an open question in systems neuroscience. Pairwise connectivity is not uniform in the neocortex (Shimono & Beggs, 2015; Perin et al., 2011; Silberberg & Markram, 2007; Song et al., 2005; Jiang et al., 2015, Karnani et al., 2016; Kwan & Dan, 2012, Cossart, 2014; Bonifazi et al., 2009). For example, it is known that area V1 pyramidal units share stronger physical and functional connections with each other if they are similarly tuned (Gilbert & Wiesel, 1989; Ko et al., 2011). However, how multi-neuronal ensembles of neurons coordinate to form computationally meaningful units remains incompletely understood.

By analyzing the patterns of firing of individual neurons during epochs of spontaneous activity (Luczak et al., 2007, Miller et al., 2014, Ringach, 2009, Carillo-Reid et al., 2015, Carillo-Reid et al., 2016) in layer 2/3 of mouse area V1 we identified pyramidal cells whose firing tends to immediately precede the firing of a neighboring interneuron with high statistical significance (Fig.2). The interneuron with its “pyramidal partners” forms an “IPP- cluster”. IPP-cluster members have higher probability to fire in synchrony with each other, even when we exclude epochs during which they might be linked via the firing of their partner interneuron (Fig.3). They are also likely to be inhibited together by the firing of their partner interneuron (Safari et al., 2017, Rikhye et al., 2017).

IPP-clusters were identified purely based on the temporal structure of individual unit firing during epochs of spontaneous activity, in the absence of visual stimulation. In spite of this, IPP-cluster pyramidal members share direction preference between themselves and with their partner interneuron (Fig.5), strongly supporting the idea that they have functional significance. This is further corroborated by the fact that pyramidal members of IPP- clusters exhibit stronger pairwise functional connectivity with each other than with members of other IPP- clusters, suggesting they form a functionally cohesive group. Co-activated pyramidal units are more efficient at transmiting “units” of information to downstream targets (Smyrnakis & Smirnakis, 2013). They also need to be inhibited together in order to unambiguously designate the end of information transmission. The ability of functionally similar IPP-cluster members to be co-modulated together suggests that IPP-cluster functions as basic computational unit, the synchrony and burst firing of which is regulated by the activation of its “partner” interneuron.

Spontaneous activity patterns suggest that the L2/3 network in area V1 conforms to a “small-world” architecture (Fig.1G-H). At the scale in which we examine the system (∼250μm), IPP-clusters can be tentatively identified as “small-worlds” interconnected by a relatively small number of “shared” units, i.e. units that overlap between distinct IPP-clusters. “Shared” cells preferentially link IPP-clusters with similar properties (Fig.5F), leading to the relative integration of similarly tuned, versus the relative segregation of orthogonally-tuned clusters. A picture then emerges according to which, under appropriate conditions, computationally relevant IPP-clusters can interact via “shared” cells to form functionally significant super-groups, yoked together for the computation of specific properties (like feature detection) or for the performance of specific logical operations.

This network scaffold of IPP clusters is not hardwired at birth but undergoes substantial refinement during postnatal development. From before eye opening to juvenile adulthood IPP-cluster size and overlap between IPP-clusters decreases, the degree of functional connectivity per IPP-cluster pyramidal member decreases, and V1 layer 2/3 network architecture increasingly evolves into a higher number of interacting but distinct “small-world” IPP-cluster neighborhoods. In agreement with this, ∼41% of IPP-cluster pyramidal members link exclusively with a single interneuron in adulthood, compared to only ∼13% before eye opening. Overall, the number of interneurons a layer 2/3 pyramidal neuron is on average linked with drops from ∼3.5 before eye opening to ∼2 in adulthood. This is consistent with prior results reporting down-scaling of pairwise connections during postnatal development (Golshani et al., 2009) especially between disparately tuned pyramidal neurons (Ko et al., 2013).

In summary, L2/3 of area V1 obeys a principle of organization where pyramidal cells are grouped together into feature-selective sub-networks (IPP-clusters or cliques (Luce & Perry, 1949)), whose firing is linked to the firing of specific, functionally similar cortical interneurons. This notion is intuitively appealing: a group of neurons representing a temporally coherent computational unit ought to be inhibited and disinhibited together, thus providing an intrinsic cortical mechanism for parsing the output of similarly tuned pyramidal cell populations to higher level areas (Roux & Buszaki, 2014, Buzsaki, 2010, Sirota & Buszaki, 2005). Since IPP-cliques appear to evolve to represent distinct stimulus features, IPP-clique interactions via “shared” (“hub”) connections may serve to improve the robustness of specific representations and help process feature conjunctions involving different aspects of the same stimulus during processes such as low-level perceptual feature-binding (Kenet et al., 2003; Gao et al., 2010; Ji et al., 2015; Miller et al., 2014, Carillo-Reid et al., 2015). Overall, our observations suggest a principle of modular organization, where IPP-clusters serve as a basic cortical modules of computation explicitly centered around individual interneurons, while computationally meaningful super-groups are formed through interactions between these basic modules mediated by shared pyramidal cells.

## METHODS

### Animals and surgery

All experimental procedures were performed in accordance with institutional and federal animal welfare guidelines, and were approved by the institutional animal care and use committee. We used C57BL/6 mice expressing Td-Tomato in Dlx5/6-positive interneurons (Madisen et al., 2010; Miyoshi et al., 2010). The mice were produced by crossing the Ai9 mice, carrying a targeted insertion into the Gt(ROSA)26Sor locus with a loxP-flanked STOP cassette preventing transcription of a CAG promoter-driven red fluorescent protein variant (tdTomato; Madisen et al., 2010), and Dlx5/6-Cre mice, carrying a Cre-recombinase gene, expressed under the Dlx5/6 promoter (The Jackson Laboratory; Miyoshi et al., 2010). Offspring mice, carrying both the flox- stopped td-tomato and Cre, express Td-Tomato in 65% of interneurons originating from the lateral geniculate eminence.

Mice were reared in a 12 h dark/light cycle until P8 – P52 and subsequently were used for recordings. During surgery, mice where anesthetized with 2% isoflurane, which was delivered in pure oxygen via tubing near the nose of the animal. Local anesthesia with lidocaine (2%) was given under the skin and on the skull. We placed a 3 mm round craniotomy above V1 and pressure injected 1mM OGB-1 with 100 μM SR-101 dissolved in Pluronic via a glass pipette 200 μm below the dura (Stosiek et al., 2003). All injection sites were located 2-3 mm lateral from midline and 1–1.5 mm frontal to the transverse sinus, placing them in visual cortex (Wang and Burkhalter, 2007). After injection, we fixed a glass coverslip above V1 with Vetbond glue. We kept eyes moisturized using a topical eye ointment (polydimethylsiloxane-200, Sigma-Aldrich). The isoflurane level during recordings was maintained at 0.6% in adult animals. For younger pups the isofluorane level was set to 1% during recording, as pups are less sensitive to isoflurane anesthesia. We collected on total data from 21 FOVs from 18 animals (6 animals per age group).

### Two-photon imaging

We used a Prairie Ultima-IV two-photon microscope with custom modifications, fed by a Chameleon Ti:sapphire Ultra-II laser and equipped with two Hamamatsu photomultiplier tubes. PrairieView software (version 4.1.1.4) was used to control the laser and collect images (Prairie Technologies). We imaged cells 120–200 microns below the pia, in layer 2/3 of mouse V1. The laser was set at a wavelength of 820 nm. At this wavelength, red fluorescing cells were either Td-Tomato expressing interneurons or SR-101-stained astrocytes. We selected an FOV containing 100–320 cells for imaging and acquired images using a 20x objective lens (0.95 NA water immersion, Olympus) at acquisition speeds ranging from 3.2 to 11 Hz, depending on FOV size. To separate interneurons from astrocytes we examined their morphology (astrocytes have distinct visible processes), shape and frequency of their calcium events, and the presence of red Td-tomato fluorescence before the injection of OGB-SR-101 mixture (when available). Most interneurons recorded from adult animals also showed strong visual responses to drifting gratings and / or textures.

### Visual stimulation

We constructed visual stimuli using the MATLAB Psychophysics Toolbox (www.psychtoolbox.org). We used drifting square-wave gratings with a temporal frequency of 2 Hz and a spatial frequency of 0.05 cycles/°. Additive plaid patterns were constructed by summing up component gratings of 50% contrast (Smith et al., 2005, Palagina et al., 2017). We used plaid patterns with the cross-angle 120° between the component gratings. Each visual presentation trial lasted 5 s: 2 s of visual stimulus presentation, followed by 3 s of spatially uniform illumination. We kept mean luminance constant throughout both the background and the stimulation periods. We presented stimuli on a flat LCD monitor (Dell), which was located 27 cm from the mouse eye, and covering 60°x80° in the contralateral monocular visual field. Presentation was monocular. We covered the nonstimulated eye with ointment and black foil during the experiment.

### Patch-clamp recordings

Whole-cell and loose-patch recordings were obtained with a Heka EPC-10 USB amplifier in current-clamp mode using standard techniques (Margrie et al., 2002). 6–8 MΏ glass pipettes filled with an intracellular solution (in mM: 105 K-gluconate, 30 KCl, 10 HEPES, 10 phosphocreatine, 4 ATP-Mg, and 0.3 GTP, adjusted to 290 mOsm and pH 7.3 with KOH, containing 10 μM Alexa Fluor-594 or tetramethylrhodamine dextran; Invitrogen) were advanced under two-photon visual guidance, initially with ∼100 mbar pressure, then ∼40 mbar when the tip reached ∼50 μm under the dura. We reduced pressure to ∼20 mbar when approaching a cell. Once resistance increased to ∼150% of the initial value, laser scanning was stopped and up to 200 mbar negative pressure was applied, until the resistance increased up to 200 MΏ. When successful, a gigaohm seal was typically formed within 2 min. The pipette was retracted carefully by a few micrometers to avoid penetration of the interior compartments of the cell during break-in. Then, ∼200 ms pulses of negative pressure (starting at 10 psi and increasing gradually) were applied via a Picospritzer with a vacuum module until the patch of membrane was broken. Fast pipette capacitance was compensated before break-in, and slow capacitance was compensated afterwards.

### Population bursts: extraction of calcium events’ onsets and definition of the avalanche-related activity

We typically recorded ∼10-40 minutes of spontaneous activity per FOV (1-4 movies). The acquired raw fluorescence movies were motion-corrected using cross-correlation between subsequent frames of the movies containing red (Td-tomato- and SR-101-based) fluorescent signals. This allowed to remove slow xy-plane drifts from the movies. After motion correction, we used ImageJ software (Abramoff et al., 2004) to draw the ROIs of cells around cell body centers, staying 1–2 pixels (∼2.4 microns) from the margin of a cell to avoid contamination from neuropil signals. We then extracted centers of mass of the ROIs, averaged the signals of cell ROI pixels and converted them to dF/F. To determine the onsets of OGB-1 spontaneous calcium responses, the dF/F timecourse for each cell was thresholded, using the noise portion of the data, to 3 standard deviations above noise. We applied moving average filter (3 s window) to the dF/F and then subtracted the result from the dF/F to obtain the noise and calculate the individual threshold for each cell. We used current-clamp recordings in 3 cells to verify that the above procedure was faithfully detecting the onsets of actual action potential bursts. We aligned the calcium event onset series with corresponding current-clamp recordings. Then, the 4 second periods after each event onset, corresponding to the rise and decay of the calcium transient, were taken out of the current clamp recording, and the remainder of it was used for baseline calculations. We then determined if any frames during and 2 seconds after event onset frame contained action potentials, and how many. The 2 second window encompasses the fast rise and peak area of calcium event, where most action potentials occur (Vogelstein et al., 2010). We determined the median spike rate in each 2 second window across all events, and determined whether any individual frame in this window contained more than 2 action potentials (doublets and multiplets). Then we broke the baseline portion of the recording into 2 second intervals, and repeated this procedure for those intervals. ∼77% of calcium events detected by this method corresponded to doublets, triplets etc. of action potentials, while single action potentials were caught in only ∼23% of cases. In contrast, the parts of the recordings that did not have detected events (baseline), consisted mainly of solitary action potentials; doublets and multiplets appeared in only 35% of baseline trials. In addition to this, the action potential packets detected by our procedure contained larger spike count than those belonging to baseline trials: median spike rate was 3.25 Hz for event trials compared to 0.5 Hz for baseline trials. This shows that our procedure discriminates in favor of cellular events related to reverberating recurrent population activity, as the events mostly contain bursts of action potentials, ocurring at higher rate, as opposed to sparcer solitary spikes observed during baseline periods.

Detected onsets then were used to form an eventogram for each FOV movie. We defined population bursts as periods of time when two or more cells had onsets of calcium responses in one or more subsequent frames. The first frame of such burst is preceded by a frame that is empty. Solitary events, even occurring in temporal proximity (e.g. separated by one empty frame) were discarded from the eventogram, so that only burst-related activity was used in further analysis.

### Small-world network parameters and analysis

Determination of ‘linked cells’ for small-world analysis was made based on presence of significant pairwise linear Pearson correlation between cells. To determine if the Pearson correlation coefficient in each pair was significant, we used simulated null distribution of Pearson coefficients obtained from circularly shuffled data. The onsets of cellular events were circularly shuffled by applying a variable offset [1<offset<(movie length-1)], where movie length is total number of frames in a given movie, for each cell. The offset was selected randomly. Shuffling was repeated 10000 times to generate the distribution of correlation coefficients that can occur by chance for each cell pair. Each pair’s null distribution’s maximum value was used as a threshold to determine if the real correlation coefficient value was significant (e.g. exceeded the threshold). Next, we examined if the cell pair’s real correlation value also was above the median correlation observed in an examined FOV across all cell pairs. If both thresholds were satisfied, the cell pair was accepted as having a significant functional link. Links obtained through this procedure were then used to calculate degree of connectivity across the network nodes (cells).

Small-world networks are defined by a relation between the clustering coefficient and the short average minimum path length. The clustering coefficient (C) for an individual cell (node) is the ratio of the actual links (r) formed between all the linked partners (k) of the cell over the number of all possible links:

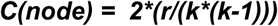

The global clustering coefficient (***Cg***) is then determined by averaging the clustering coefficients of all nodes. Small-world networks also have short path lengths (***L***), also a common property of random networks. Path length is a measure of the distance between nodes in the network, calculated as the mean of the shortest geodesic distances (number of edges) between all possible node pairs. For nodes *i, j, **L*** is determined by:

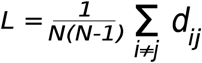

where *dij* is the shortest geodesic distance between nodes (i,j) and *N* is the number of the cells in the network. To calculate the small-world factor, we compare the global clustering coefficient and path length of the actual network with the global clustering coefficient (***Cg,ran***) and path length (***Lran***) of a random network with the same number of nodes and same mean number of links per cell. Such a network was created by randomly redistributing the existing links in the actual network across the nodes. Then we calculated the small world factor (SMW), as follows:

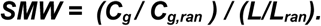

SMW > 1 is used as a criterion to classify the network as a “small-world” net (Humphries et al., 2006, Sporns, 2011), since it argues for high average clustering coefficient relative to path length compared to otherwise equivalent randomly connected nets. About 8% of nodes in the network had limited connectivity: they either were not connected to any other nodes in the network, or were connected to only to a very restricted subset of nodes, which resulted in their individual average path length value to be close to zero. We checked if their inclusion in the calculation affected the resulting SMWs. To do so, we either imputed the mean path length for them with the mean path length across the connected nodes, or excluded them from the calculation all together. In both cases the SMWs decreased only slightly, with lowest values staying above 1.64 in all cases.

### Identifying “interneuron pyramidal partners” (IPPs)

**T**o determine if the activation of pyramidal cells and interneurons in the context of multineuronal population bursts was random, or alternatively, groups of specific pyramidal cells and interneurons were working in concert, we compared the probability of individual pyramidal cell’s having a calcium event 600 ms before and 600 ms after the event in the interneuron. For each pyramidal cell (PC), we generated a probability difference measure, by subtracting the probability of PC event 600 ms before the interneuronal event from the probability of PC event 600 ms after the interneuronal event, and selected out the cells with positive difference values for further analysis, as these cells potentially had higher probability to be activated shortly before interneuronal events. The probability difference measure was calculated over (13-92) instances of inteneuronal activations. If the interneuron had less than 12 in-burst calcium events, we did not consider it for further analysis.

We then generated 10000 instances of surrogate data sets by temporally shuffling the event onsets in these potential PC partners, and leaving the interneuronal event onsets intact. For each shuffled data set and each potential PC partner, a surrogate ‘chance’ probability of a pyramidal cell to be activated after the interneuronal event was generated (10000 points). Next, for each tested PC, the real probability to have an event before the interneuronal event was compared against each ‘chance’ probability point in this null distribution, and the fraction of instances when the chance probability was equal or exceeded the real probability was used as a p-value for each interneuron – pyramidal cell pair. Individual cells’ p-values were than corrected for multiple comparisons (by multiplying them by the number of tested PC partners), and pyramidal cells with corrected p-values below 0.5 were selected as the partners of the interneuron. 95% of adjusted p-values were below 0.044, with median value at 0.016.

The groups of pyramidal cells whose probability of firing was significantly increased before the event in the interneuron, were then called “interneuron pyramidal partners” (IPP) and assigned to a single cluster (IPP- cluster).

The 600 ms window was selected based on the latencies of responses of oriented interneurons and pyramidal cells in response to preferred stimulus in 4 FOVs (from 4 adult animals, calcium data), and the dynamics of ongoing excitatory and inhibitory population inputs into the layer 2/3 pyramidal cells (intracellular recordings data), obtained from Douglas & Martin (Douglas & Martin, 1991). In individual layer 2/3 neurons, ongoing network activity resulting from visual stimulation causes fast population EPSP, followed by prolonged population IPSP, which lasts for ∼ 250-350 ms after the population EPSP onset (Douglas & Martin, 1991). In our calcium data recorded under preferred visual stimulus there was ∼140 ms median difference between interneuronal and pyramidal cell population latency (calculated across all trials and all cells in the given FOV). For the trials where the pyramidal cell’s responses had shorter or equal latency compared to interneuronal responses, median latency difference between interneuronal and pyramidal populations was ∼600 ms.

### Global functional connectivity of the pyramidal cells and group correlation strength of the IPP clusters

We next checked if event trains of individual IPP cluster members had different group correlation strength between themselves compared to other pyramidal neurons in the FOV, that did not belong to the cluster. For this, we first determined which pyramidal cells pairs in the FOV had significant cross-correlations within the test window used to determine the membership of pyramidal cells in IPP clusters. The cross-correlation analysis was done on the eventogram binary data within a [-600 ms 600 ms] window, where t=0 corresponds to the pyramidal cell’s event. Pairwise linear Pearson coefficient was computed for every PC pair. To determine if the coefficients were significant, we generated 10000-iteration null distribution of values, by circularly shuffling the event onsets in every cell. The displacement values for each cell were selected at random. Using this 10000-point null distribution, we looked if the actually observed cross-correlation for a specific pair was outside the 99.7% values of the null distribution to accept the value as significant and not occurring by chance. Pairs with a significant value were counted as a functional ‘link’. The non-significant coefficients were set to zero. We then used the resulting matrix of significant functional links (’significance matrix’) to explore the group correlation strength within the IPP clusters.

For this, we first removed periods of interneuron activation ([-600 ms, 600 ms] perios around each interneuronal event) from the FOV’s eventogram. Next, the pairwise cross-correlation was computed between every pyramidal cell pair in the remaining portion of the eventogram, using a window of [-600ms 600ms]. We then used significance matrix to determine if the link between a particular pair of cells was significant. Non-significant links’ correlation values were put to zero. Then, mean cross-correlation coefficient was measured for PC pairs of the same IPP cluster. To determine if this mean peak cross-correlation was significantly different from control, we generated 10000 surrogate cell groups, size matched to the tested IPP cluster, by randomly selecting PCs out of the FOV. We used the following criteria to select the member cells for these control groups (see also Fig.3):

1. Their pairwise distances to the partner interneuron were restricted to match the pairwise distance range of the tested IPP cluster members to the partner interneuron. For this, we averaged distances from interneuron to each partner pyramidal cell in the cluster. The distance envelope was set to mean±3 s.d. from the interneuron.
2. The cells did not belong to an IPP cluster.
3. We then calculated the mean pairwise cross-correlation coefficient of the eventogram between the members of the surrogate group and the members of the tested IPP cluster. This was repeated 10000 times to arrive at the null distribution of mean correlation strength of randomly selected cell groups in the given FOV. Using this null distribution, we determined if the IPP-cluster’s mean pairwise cross-correlation peak value was outside of P=0.997 interval (3 standard deviations from the mean).

### Global functional connectivity of the pyramidal cells during multineuronal bursts

We next used the significance matrix to explore the refinement of functional connectivity in the course of early development. We set all non-zero values to one, denoting a presence of a functional link between the cells, while non-linked cellls were assigned a zero. We next used the resulting matrix to determine the amount of links made by pyramidal neuron that participated in a singular IPP cluster (’exclusive’ cell) versus the amount of links made by pyramidal neuron participating in more than one IPP cluster (’shared’ cell).

### Selectivity for the direction(s) of stimulus motion

The dF/F of each evoked calcium data was deconvolved using the algorhithm of Vogestein et al. (Vogestein et al., 2009). Resulting output data were used to construct cell’s tuning curve for drifting gratings and moving plaids. We evaluated *t*he selectivity of each cell for the direction of motion of drifting grating using the direction selectivity index (DSI):

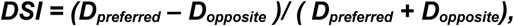

where *D_preferred_* is the response for motion to the cell’s preferred direction, and *D_opposite_* to the opposite direction. Highly selective cells have DSI near 1, while a 3:1 response difference would result in a DSI of 0.5. The cells that generate bi-peaked tuning curves for the drifting grating are characterized by orientation-selectivity index, as those peaks are typically opposite or near-opposite each other:

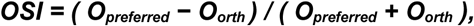

where *O_preferred_* is the direction axis (orientation) along which peak responses occurs, while *O_orth_* is the direction axis (orientation) orthogonal to peak response axis. We considered the cell to be tuned for direction and/or orientation in case that either of the metrics exceeded 0.45. Out of 23 visually-responsive interneurons in adult set (collected from 4 FOVs), 15 were tuned for the direction and / or orientation of the moving gratings (DSI or OSI equal or above 0.45). We than examined the tuning curve of each interneuron and excluded 3 cells with exceptionally broad tuning, that had more than 3 significant peaks in their tuning curve (for this we determined the amplitude of the largest peak, and accepted any other peaks that were above half of that amplitude as significant). This left us with 12 interneurons for the grating condition.

For all examined tuned interneurons we first identified the dominant peak in their tuning curve. Most cells also had additional smaller peaks in their tuning curves, typically located opposite of dominant peak, as most of the oriented neurons in V1 show a degree of orientation preference. We thus looked if the interneuron had additional peaks located in the 120 degrees window centered on the direction opposite to the dominant peak. The minor peak was considered significant if the amplitude of the local maximum was signifcantly larger than zero. We considered both peaks as ‘preferred direction’ when evaluating the alignment of tuning between the tuning curves of the interneuron and its partner pyramidal cell (see Fig.5).

For plaid condition, we again first selected the tuned interneurons based of direction-selectivity of interneuron’s responses. First, we calculated the direction-selectivity index similar to grating case:

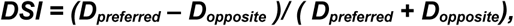

*where D_preferred_ is the response for motion to the cell’s preferred direction, and D_opposite_ to the opposite direction.* Highly selective monopeaked cells have DSI near 1, while a 3:1 response difference would result in a DSI of 0.5. However, most cells in V1 generate double-peaked responses for moving plaids, for two opposite directions of motion (Palagina et al., 2017, Muir et al., 2015). To detect these cells, we used plaid direction-selectivity index:

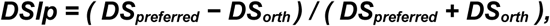

*where DS_preferred_* is the sum of responses for motion to the cell’s preferred directions, and *DS_orth_* is the sum of the responses to the orthogonal directions. If either of these indexes reached 0.45, the cell was accepted as tuned for the plaid’s direction(s) of motion. Next, we removed the broadly tuned or multipeaked interneurons, and determined the locations of dominant directions similarly to the grating case. Thus, for plaid condition we used 10 tuned interneurons.

We then examined the alignement between the tuning curves of the interneurons and the pyramidal cells found in their corresponding clusters. First, we examined the population tuning curves of pyramidal cluster members vs interneuron (Fig.5C). Since the communal tuning curve appears to be dominated by oriented pyramidal members, we proceeded to describe the relationshipe between their tuning and the tuning of the interneuron. For each cluster, we selected out cells that had sufficient direction or orientation tuning. First, we removed cells whose OSI or DSI was below 0.45. We examined the tuning curves of the remaining cells and removed cells with noisy peak responses and cells whose tuning curves had more than two significant peaks (a significant peak reached 2/3 of the amplitude of the dominant peak). For the remaining cluster members, we determined the smallest offset of the preferred direction of each pyramidal neuron from the preferred directions of the partner interneuron.

### Statistical tests

We used Wilcoxon ranksum test (WRS) to test for group differences, and Wilcoxon signed rank test (SR) in cases when the group data was paired. All points in the errobar plots are shown as mean±sem, unless otherwise noted.

---

**Extended Data Figure 1.**
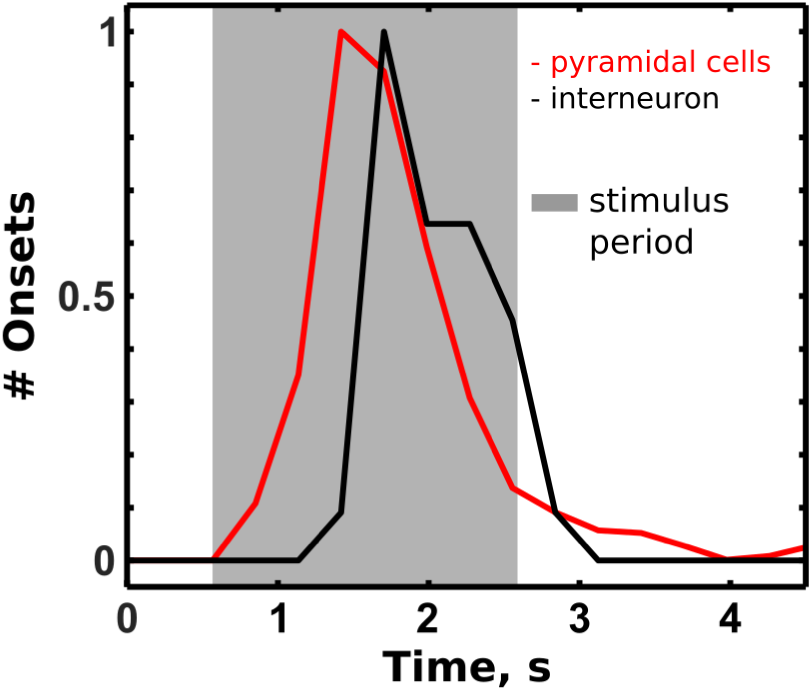
Relationship between interneuronal and pyramidal cell activity as measured by calcium imaging. To compare the latencies of interneurons and pyramidal cells, we selected first cells that displayed orientation or direction selectivity. For each cell, we singled out trials when the cell was driven by preferred stimulus. For each trial, the poststimulus latency was determined. Next, latencies were grouped across the iterneurons and pyramidal cells to be compared to each other. **Red:** distribution of onsets of calcium responses of all oriented pyramidal cells found in the FOV (1554 trials from 78 cells, cells are driven by the grating of preferred direction). **Black:** distribution of onsets of calcium responses of single oriented interneuron from the FOV (20 trials, cell is driven by the grating of preferred direction). The distributions are normalized by peak values. Interneurons and pyramidal cells come from the same FOV. Gray area: period of stimulation with the grating of preferred orientation. Frame period for this FOV was 0.284 s. The medians of the distributions are separated by 284 ms. The median lag between pyramidal cells and interneurons across 4 FOVs was ∼140 ms when all pyramidal cell trials were considered. However, if we only considered pyramidal cell trials with shorter latencies relative to interneuronal trials, the lag increased to ∼ 600 ms, which was subsequently chosen as a window for determining the presence of functional link between pyramidal cell and interneuron.

**Extended Data Figure 2.**
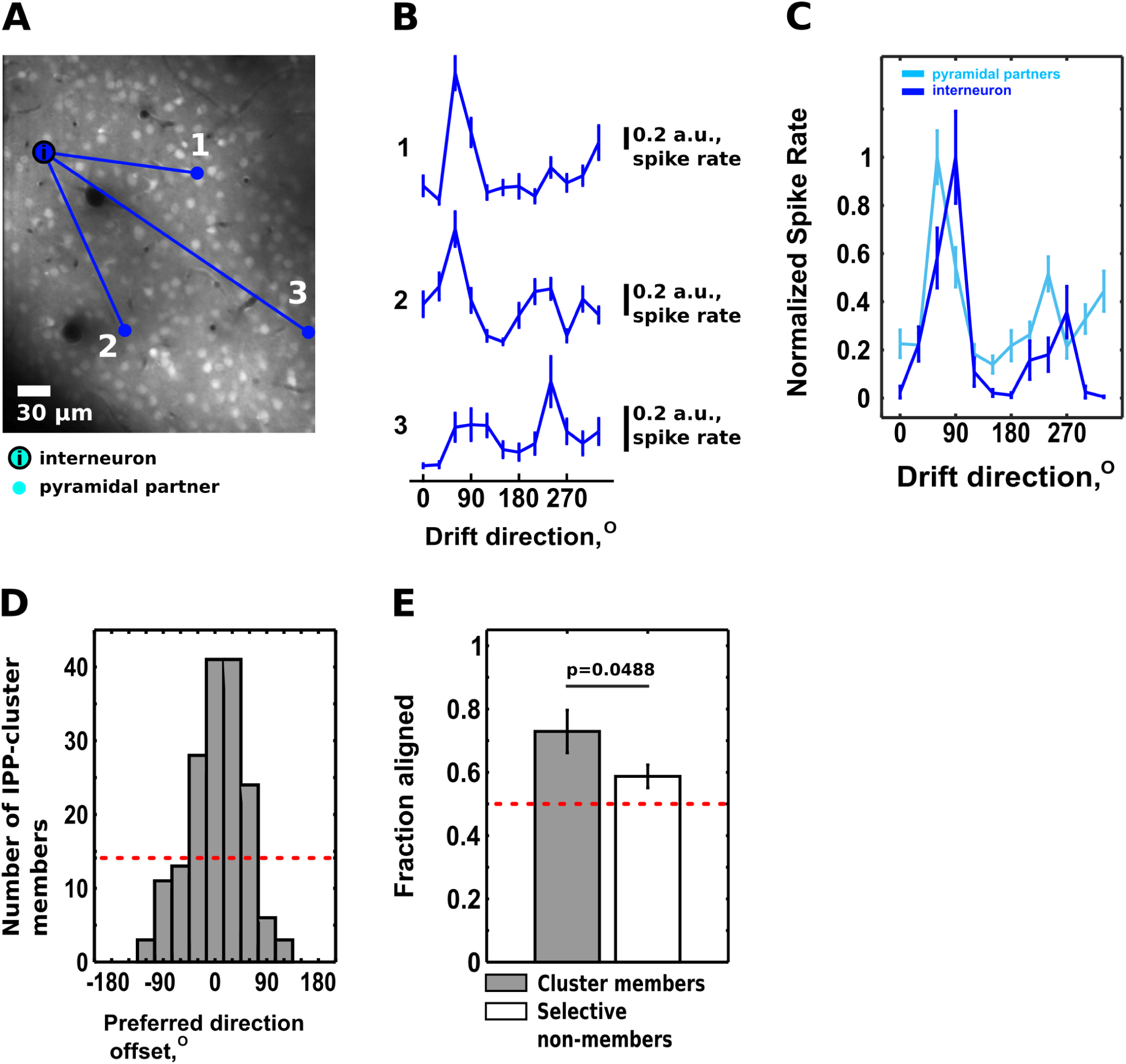
Functional properties of IPP-clusters tested with moving plaids (cross-angle: 120°). **A**. Example IPP-cluster with 3 pyramidal cell members. Partner interneuron is marked with “i”. Numbers denote pyramidal cells whose tuning is shown in B. B. Tuning curves of the cells numbered in A, when stimulated with moving plaids (with respect to the pattern direction of the plaid). Vertical bars: spike rates inferred from deconvolved dF/F (arbitrary units). **C.** The population tuning of IPP-cluster members (cyan) and the tuning of the partner interneuron (blue) share high similarity. Median linear Pearson’s correlation between the tuning curves for the partner interneuron and its IPP-cluster members was ∼0.508 (n = 10 IPP-clusters). **D.** Distribution of differences between the preferred direction(s) of tuned interneurons and corresponding IPP-cluster’s pyramidal members. When stimulated with plaids, tuned interneurons typically have a bi-peaked direction tuning curve (see panel C, blue tuning curve). The majority of tuned IPP cluster members have the same preferred direction as the partner interneuron, when stimulated with moving plaids (∼73% of tuned IPP cluster members have their preferred direction within ±45° of interneuron’s preferred direction(s)). **E**. <u>***Gray Bar:***</u> Fraction of IPP-cluster pyramidal members whose preferred direction is aligned (to within ±45°) with that of their partner interneuron. <u>***White Bar:***</u> Fraction of tuned pyramidal cells that do not belong to the specific interneuron’s cluster (n = 10 IPP clusters, p = 0.0488, Wilcoxon ranksum test). <u>***Red dotted line:***</u> fraction of cells expected to be aligned by chance. Data are expressed as mean ± sem, tuned cluster members: 0.73±0.07, tuned non-members 0.59±0.04.

**Extended Data Figure 3.**
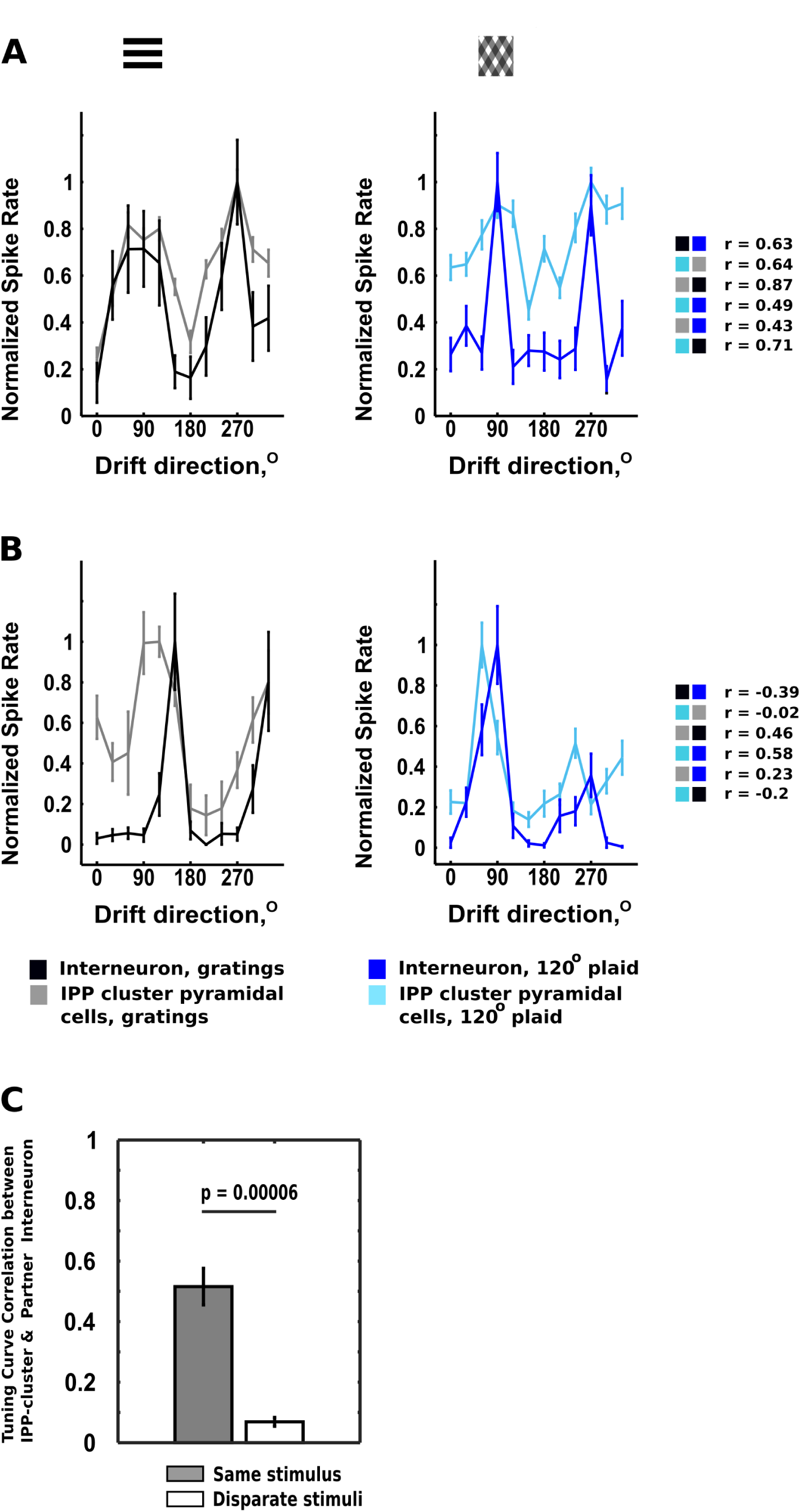
The similarity of functional properties between IPP-cluster members and their partner interneuron persists across different stimuli. **A,B.** We examined how the tuning of the cluster members and their partner interneuron changes with the change in the visual stimulus. We selected interneurons that showed visual stimulus-driven responses to 2 different moving stimuli: gratings and wide (120° cross-angle (CA)) plaids, and showed significant direction or orientation tuning in response to at least one type of stimulus. In mouse V1 oriented cells typically generate responses to both plaids and gratings; however, the peaks of the cell’s tuning curve often shift between the two stimuli (Palagina et al., 2017; Juavinett & Callaway, 2015; Muir et al., 2015). The population tuning of the IPP-cluster members closely followed the tuning of the partner interneuron, even if interneuron’s tuning properties and peak responses shifted massively between different stimulus types. Left panel: tuning curves generated by the interneuron and its partner IPP cluster for gratings. Right panel: tuning curves generated by the interneuron and its partner IPP cluster for plaids. **A.** Example of an interneuron with stimulus-invariant direction tuning and its partner cluster. The interneuron displays bimodal tuning curve both when stimulated with moving grating (left panel) and moving plaid of 120° cross-angle (120°-CA, right panel). Preferred directions of the interneuron for the 120°-CA plaid are aligned with the preferred directions under the grating condition. The population tuning curve derived from the IPP-cluster pyramidal members follows the tuning of the interneuron closely for both grating and plaid conditions. On the right we list linear Pearson correlation coefficients between the different pairs of tuning curves (color-coded). Population tuning curves of IPP-cluster members have a high degree of similarity (r=0.49 and r=0.87) to the tuning curve of their partner interneuron, irrespective of stimulus. Interstimulus correlations are also high for both interneuron and the cluster (r = 0.43−0.71) due to the invariance in diection-tuning. **B.** An example of the interneuron whose direction tuning changes considerably between moving grating (left panel) and moving plaid (right panel) stimuli. The majority of oriented interneurons and pyramidal cells in V1 showed such change in direction preference. In the example shown, the peaks of the tuning curve for the grating are shifted by ∼60° relative to the peaks of the tuning curve for the plaid. Strikingly, the population tuning of the IPP cluster closely follows the population tuning of the interneuron between the conditions: the IPP- cluster’s tuning curves remain aligned with the interneuron’s tuning curve even when the stimulus and interneuron’s responses change drastically. In this case interneuron’s preferred directions for plaid and grating are 60° apart, but the Pearson correlation between the tuning curves of the interneuron and its IPP cluster remains high: 0.46 for the grating condition and 0.58 for the plaid condition. In contrast, interstimulus correlations are much lower for both interneuron and IPP-cluster: r = −0.39 – 0.23. **C.** As expected, Pearson correlation coefficient between (i) IPP-cluster population tuning curves and (ii) partner interneuron tuning curves is much higher for the same versus across different visual stimuli (same stimulus: n=22 pairs, disparate stimuli: n=22 pairs, p = 0.000062, Wilcoxon ranksum test).

**Extended Data Figure 4.**
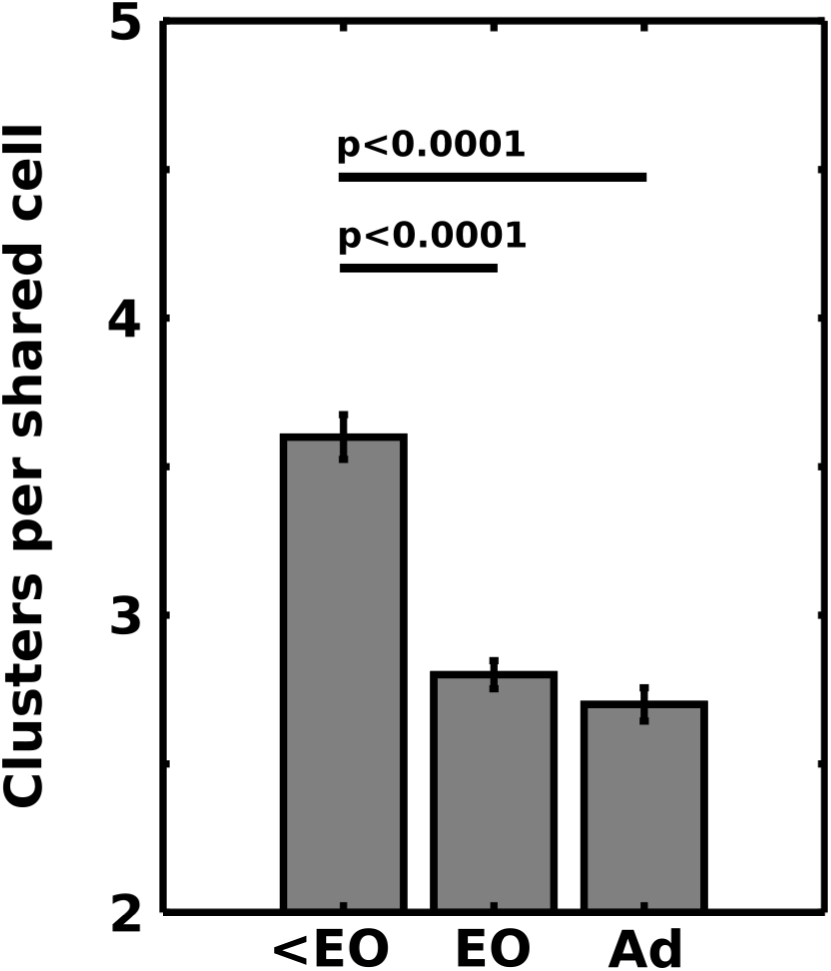
The number of IPP-clusters that a particular “shared” (“hub”) cell participates in decreases over postnatal development from 3.7±0.075 (<EO, n = 428 shared cells, 7 FOVs) to 2.8±0.047 (EO, n = 401 shared cells, 7 FOVs) to 2.7±0.056 IPP-clusters / shared cell (Adult, n = 243 shared cells, 7 FOVs).

